# cpCST: A New Continuous Performance Test for High-Precision Assessment of Attention Across the Lifespan

**DOI:** 10.1101/2025.06.22.660941

**Authors:** MacKay-Brandt, Garcia-Barnett, Gan, Ripley, Gazes, Milham, Colcombe

**Author notes:** **Correspondence:** Stan Colcombe.

## Abstract

Assessing sustained attention presents methodological challenges, particularly when spanning diverse populations whose baseline sensorimotor functioning may vary significantly. This study introduces the Continuous Performance Critical Stability Task (cpCST), a novel paradigm combining high-density sampling of behavior (30Hz), individualized calibration, and fixed-difficulty assessment to measure attentional control. In a sample of 166 adults (ages 18-76), we evaluated the psychometric properties of the cpCST’s instantaneous reaction time (iRT) metric derived through dynamic time warping. Results demonstrated exceptional reliability (bootstrap split-half r = .999), age invariance, and predictive validity for cognitive performance (flanker and Woodcock-Johnson) and cardiorespiratory fitness (est. VO2max). The cpCST achieved high temporal efficiency, with just two minutes of data correlating at r = .94 with full-task performance, outperforming a standard arrow-based flanker task. The cpCST’s individualized calibration effectively isolated attentional control processes from baseline sensorimotor function, eliminating age-related slowing effects typically observed in reaction time tasks. This approach offers methodological advantages for lifespan studies, clinical populations, integration with neurophysiological measures, and computational modeling approaches while addressing limitations of existing attention assessment paradigms.

## 1 Introduction

Attention is an intuitive concept that is considered a core component of cognition and everyday functioning (Baddeley, 1996; Duncan, 1986; Norman & Shallice, 1986; Posner & Petersen, 1990), exhibiting a clear trajectory of early life maturation and later-life decline (Fortenbaugh et al., 2015; Tipper et al., 1989). Across the lifespan, attentional processes are linked to the successful navigation of a host of everyday behaviors (Barriga et al., 2002; Bogdanova et al., 2016; Gross, 2015; Halperin, 1991; Kinsella, 1998; Racer & Dishion, 2012; Stierwalt & Murray, 2002); and like many apparently simple behaviors, it can be challenging to define and measure (Anderson, 2021; Shi et al., 2019; Unsworth et al., 2024; von Bastian et al., 2020; Yangüez et al., 2024).

Experimental and clinical work focused on the measurement of sustained attention has produced a wide selection of continuous performance tasks (C. K. Conners et al., 2003; Cooper et al., 2017; Dinges & Powell, 1985; Gronwall, 1977; Homack & Riccio, 2006; Klee & Garfinkel, 1983; Leark et al., 2007; Lopez-Garcia et al., 2016; Mackworth, 1948; Mirsky & Rosvold, 1963; Robertson et al., 1997; Sahakian & Owen, 1992; Schmidt et al., 2024; Servan-Schreiber et al., 1996). These attention tasks are tuned to capture behavioral features thought to contribute to successful performance or identify specific areas of deficit, based on the paradigm and study population of interest.

As outlined below, opportunities exist to augment or improve upon existing paradigms through novel behavioral sampling, dynamically adaptive assessment, and task calibration approaches - amongst others. Here we describe the Continuous Performance Critical Stability Task (cpCST), which builds upon an established sensorimotor integration task to create a novel attention task featuring high-density behavioral sampling, dynamic adaptation, and effective behavioral calibration across the lifespan. We present an initial psychometric evaluation of the cpCST’s primary outcome metric (instantaneous reaction time; iRT). We also examine predictive validity of the cpCST iRT to flanker task performance, Woodcock-Johnson Intellectual Ability and Achievement scores, as well as a measure of cardiorespiratory fitness (VO2max), before discussing the advantages of the cpCST paradigm in relating physiological and brain timeseries to participant behavior.

### Sparse sampling is typical in continuous performance tests

In most continuous performance tests, attention is probed via button press responses at discrete intervals ranging from roughly one to several (10+) seconds apart c.f.(DiFrancesco et al., 2019; Dinges & Powell, 1985; Homack & Riccio, 2006); attentional lapses are inferred on the occasion of delayed, missed, or incorrect responses. Despite the relatively sparse sampling of behavior (<1Hz - once every second or longer), these response time studies have demonstrated attentional fluctuations over time (Castellanos et al., 2005; Decker et al., 2023; Di Martino et al., 2008; Esterman et al., 2013; Jackson & Balota, 2012; Smallwood et al., 2004); however, higher density sampling of behavior may more effectively characterize the maintenance of focus over time, moment-to-moment fluctuations, and/or lapses in attention.

### Altering task features in cognitive tasks

Administering reaction time tasks to older adults, children, or clinical populations often requires adjustments to various parameters such as stimulus type, stimulus modality (audio vs. visual), presentation and response durations, response interval, interstimulus intervals, stimulus set sizes, or proportion of trial types (see (Cowan et al., 2010; Craik, 1986; de Souza Almeida et al., 2021; Reuter-Lorenz & Cappell, 2008; Rueda et al., 2004)). These adjustments are motivated by group differences in sensorimotor speed, working memory, auditory or visual acuity, etc. (Cerella, 1990; Conway et al., 2003; Denckla, 1996; Jacobson et al., 2011; Kail, 1991; Owsley, 2016; Salthouse, 1996; Verhaeghen & Cerella, 2008; Wingfield et al., 2005). While accommodations such as these allow for versions of standard neurocognitive tasks to be applied across a wider range of individuals, they raise concerns regarding the comparability of results across test variants (Best & Miller, 2010; Cooper et al., 2017; Hedge et al., 2018; Parsons et al., 2019). Additionally, these changes are applied under the assumption that the altered parameters are uniformly appropriate to the group in question (e.g. trading arrow shapes for cartoon fishes in the ANT-C task or an increase in presentation duration for older adults (Rueda et al., 2004)), despite well-documented heterogeneity within groups (Fair et al., 2012; Lindenberger & Baltes, 1997; S. Logan et al., 2023).

### Fully adaptive tasks

Rather than assuming that a single set of task adaptations will be equally appropriate across a given group (e.g. older adults or children), fully adaptive paradigms individualize task parameters for each participant by dynamically altering key task features in response to ongoing task behavior. Some tasks are explicitly designed to be adaptive (e.g. Stop-Signal Reaction Time (G. D. Logan & Cowan, 1984)). More recent approaches overlay adaptive procedures that alter task features such as presentation time, response windows, or set size, in response to participant performance in real time as the task evolves (Barbey et al., 2022; Draheim et al., 2021, 2024; Schneiders et al., 2011). As such, each participant’s task is custom tailored to their individual performance on that task through approaches such as staircase or Bayesian-based adaptive algorithms (Farahbakhsh et al., 2019; Manning et al., 2018). These approaches are more efficient (Attarha et al., 2024; Davis et al., 2002; Gibbons et al., 2024; Sorrel et al., 2020), and can be leveraged not only in assessment, but also training protocols (e.g.(Roheger et al., 2020)). However, they also suffer from drawbacks such as edge case and small sample size failures, induction of artifactual oscillatory ‘yo-yo’ patterns in difficulty, as well as the additional complexity involved in dynamically adapting task parameters in real time (García-Pérez, 2011; Kontsevich & Tyler, 1999; Treutwein, 1995),

### Hybrid calibrated+fixed difficulty tasks

One promising approach is to leverage the best of both dynamic adaptive approaches and fixed stable approaches. Participant ability is assessed via adaptive staircase or Baysian methods in a calibration phase. During the test phase, difficulty is set to a fixed level matching the participant’s individual ability (e.g. the stimulus onset asynchrony that resulted in > 70% accuracy) (Chen et al., 2017; Fleming et al., 2010; Lindfield et al., 1994). Under this approach, difficulty is individually tuned - thus avoiding assumptions about the appropriateness or comparability of group-specific stimulus changes. And the difficulty is fixed during the testing phase, reducing computational complexity and artefactual issues such as induced oscillatory behavior or algorithmic failure.

### The Continuous Performance Critical Stability Task

(cpCST) was designed to capture attentional dynamics from moment to moment using a simple, relatively short duration, high information density, individualized difficulty approach in order to maximize the detection of attentional control performance differences across an adult lifespan sample. The cpCST uses the hybrid-calibrated approach to first assess visuomotor ability and then fix the difficulty of a visuomotor continuous performance task based on the individual’s motor stability threshold (MST; see below). The cpCST additionally possesses useful features such as simple task instruction and continuous sampling of behavior (@30Hz; i.e. 30 times per second), providing a robust complement to existing neurocognitive tools.

The cpCST was inspired by the psychomotor tracking task developed for NASA by Jex and colleagues to evaluate pilot performance under unstable control conditions (Jex et al., 1966). This paradigm has subsequently been adopted by human and non-human primate laboratories to develop and refine human-machine interfaces (Quick et al., 2018) and understand the neural mechanisms of sensorimotor coordination (Sadeghi et al., 2024). See (Huang et al. 2005) for a similar paradigm that models event-related EEG brain dynamics. The Critical Stability Task (CST) also provides a strong foundation upon which to build a continuous self-calibrated task to assess dynamic fluctuations in attentional control. Specifically, we employed a variant of the original CST to serve as a calibration phase that established a participant’s individual MST. We then used that individualized score to set the difficulty level for that participant’s <u>continuous performance phase</u>. The goal for this new continuous performance phase was to maintain attention on a relatively easy task (set as 30% of MST, based on internal pilot testing) for a 10-minute period to capture attentional drift and the latency to respond to a drifting stimulus. See Figure 1.

**Figure 1.**
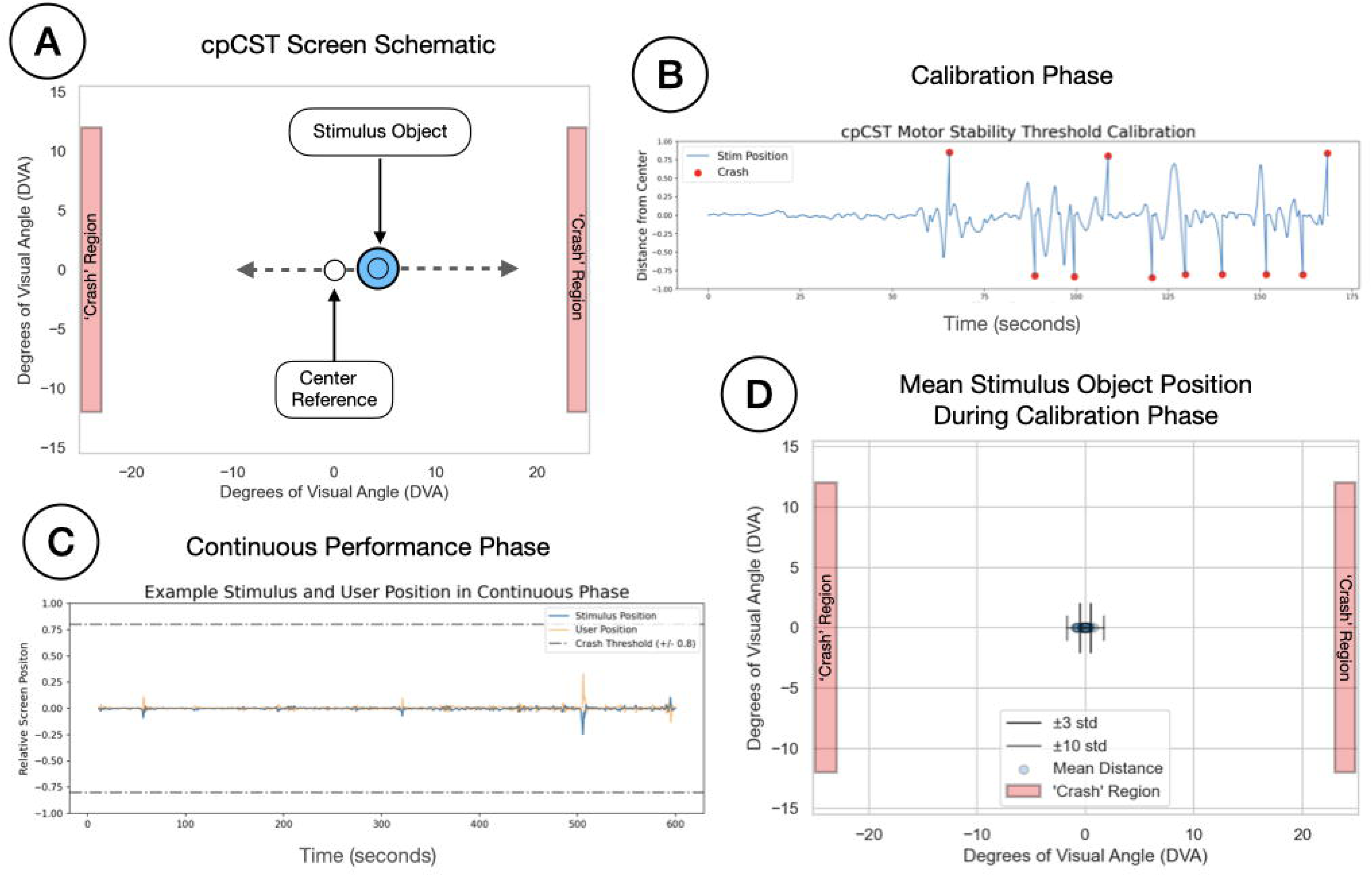
Panel A provides a schematic for the cpCST task screen. The Stimulus Object (SO) is a circle that can move in the x (left-right) dimension; participants are asked to maintain the position of the central stimulus at the center of the screen by tilting a custom accelerometer-enabled button box. The screen also contains a central reference point to provide participants with a spatial anchor during task performance, and a pair of crash region rectangles marking the out of bounds point for the SO. Panel B shows a timeseries from the SO position during a cpCST calibration phase, with time (seconds) on the x-axis and stimulus eccentricity on the y-axis. In the calibration phase, participants attempted to keep the SO at the center of the screen as difficulty (lambda) linearly increased until the participant lost control. Panel B shows the trace of an SO during the calibration phase (blue line), as well as crashes (red dots), where the SO eccentricity exceeded +/- 80% of the distance from the center to the edge of the screen. Participants perform the task through 10 crashes, and the difficulty at the time of the crash is recorded. The user’s motor stability threshold (MST) is calculated by the mean lambda value for the last 3 crashes. The MST value is then used to set the difficulty for the continuous performance phase. Panel C shows the path of the SO (blue line) and the participant tracking position (orange line) when controlling the SO during the continuous phase. Dashed grey lines show the +/- 80% crash boundary threshold. Panel D shows the mean stimulus object position during the continuous phase for all participants, as well as the +/- 3 and +/- 10 std lines. The difficulty for the continuous phase is set to 30% of the user’s MST. Figure 1. Alt Text: Four-panel figure labeled A–D. **A.** A top-down schematic of the cpCST screen: a central blue circle (“Stimulus Object”) sits on a horizontal dashed line with arrows pointing left and right toward tall red bars labeled “Crash Region.” A smaller white circle to its left marks the “Center Reference.” Axes span –25 to +25° visual angle horizontally and –15 to +15° vertically. **B.** “cpCST Motor Stability Threshold Calibration”: a blue line shows stimulus distance from centre over ∼0–175 s on a time-series plot; red dots mark each crash event. **C.** “Example Stimulus and User Position in Continuous Phase”: overlapping z-transformed traces of stimulus (blue) and user (orange) positions over 0–600 s, bounded by grey dash-dot lines at ±0.8 to indicate the crash threshold. **D.** “Mean Stimulus Object Position During Calibration Phase”: blue circles clustered near 0° on the horizontal axis show each subject’s mean position; solid whiskers extend ±3 SD and faint whiskers ±10 SD; red shaded bars at ±22° mark the crash regions.

To evaluate this novel task, we embedded the cpCST within the Nathan Kline Institute Rockland Sample II (NKI-RS2), a large-scale, community-based lifespan study. The NKI-RS2 was designed to support the development and validation of next-generation tools for phenotyping normative brain-behavioral associations and investigate the underlying neural and physiological mechanisms that promote mental health across the lifespan. This context offered an opportunity to examine individual differences in attentional control across a wide age range using a task that prioritizes continuous behavioral sampling, individualized calibration, and high-density data collection. We characterized cpCST performance in relation to broader indices of cognitive function and health. In this preliminary analysis, we report behavioral data from the cpCST from a subset of participants to describe the development of a key task performance metric (instantaneous reaction time; iRT) and establish its reliability and preliminary predictive validity on cognitive and physiological indices.

## 2 Materials and Methods

### 2.1 Participants

Participants were recruited into the Nathan Kline Institute Rockland Sample II study (NKI-RS2) through prior participation in the NKI-RS research program (Nooner et al., 2012), community outreach, and word-of-mouth. The lifespan sample recruited participants from age 9 to 75 years who were residents of Rockland, Orange, Bergen, or Westchester counties in the north suburban New York City area. All were fluent in English and had no severe physical or sensory limitations, contraindications for MRI or cardiovascular fitness testing, or acute psychiatric symptoms. Participants were excluded if they had a history of schizophrenia, schizoaffective disorder, autism spectrum disorder, or serious neurological conditions (e.g., Parkinson’s disease, traumatic brain injury, dementia). Current psychotropic medication use and serious medical conditions or metabolic disorders affecting the central nervous system (e.g., malignancy, HIV) were also exclusionary. For this preliminary analysis, we included a subset of 166 participants aged 18 to 76 years (M = 51.61, SD = 16.36), 66% female, with complete and quality controlled data for the cpCST and cardiorespiratory fitness procedure.

### 2.2 Procedures

Sample characterization data were collected via remotely administered surveys on the MindLogger Platform (Klein et al., 2020) and in-person testing. The Continuous Performance Critical Stability Task (cpCST) was administered in a dedicated testing room at the Center for Biomedical Imaging and Neuromodulation (CBIN) within the Nathan Kline Institute (NKI). The study was approved by the NKI Institutional Review Board, and all participants provided informed consent before undergoing any procedures.

### 2.3 Measures

#### 2.3.1 Continuous Performance Critical Stability Task (cpCST)

Participants were seated in front of a 61 x 36 cm computer monitor, at a distance of 65 cm. The monitor displayed a circular stimulus at the center of the screen subtending 3.17 degrees of visual angle (DVA). Screen resolution was 1920 x 1080 pixels. They were instructed to maintain the stimulus position at the center of the screen by tilting a custom-built handheld accelerometer device to the left or right to control the central stimulus speed and direction of movement. Participants were given a brief (∼2min) practice round in which they gained familiarity with the controls at a very low difficulty level prior to beginning the calibration phase.

The stimulus position was linked to an increasingly unstable system that introduced small to moderate spatial drift along a single dimension (left-right). The system’s instability was modulated in a closed-loop circuit, with instability characterized by a lambda (λ) parameter representing the gain between user input and the magnitude of the response of the system (Jex et al., 1966). This parameter sets the effective difficulty of the task. On each trial the goal was to maintain the position of the circular stimulus over a fixed central dot. This was achieved by countering the movement of the stimulus (e.g. tilting the device to the left if the stimulus drifted to the right). If the participant failed to maintain the stimulus within a predefined spatial boundary (80% of distance from the center, or +/- 22.28 DVA from center), it resulted in a "crash" and the circular stimulus was reset to the center of the screen. The position of the central stimulus and the user’s tracking position were continuously recorded at a sampling rate of 30Hz. See Figure 1, panel A.

The Continuous Performance Critical Stability Task (cpCST) employed a hybrid approach consisting of an initial calibration phase that estimated the participant’s motor stability threshold, followed by a continuous performance phase in which they performed the task at a fixed difficulty level.

##### 2.3.1.1 Calibration Phase

The calibration phase was similar to the original Jex et al. (1966) approach. Specifically, we employed a maximal performance to failure protocol similar to working memory tasks like digit span, and Corsi blocks (Corsi, 1972; Milner, 1971), and conceptually similar to the testing-the-limits approach (Kliegl et al., 1986).

During the calibration phase, participants attempted to maintain the central position of the stimulus by adaptively tilting the accelerometer device. Task difficulty (lambda) was linearly increased over time until the participant failed to control the stimulus - defined by the stimulus exceeding 80% of the distance from the center of the screen (crashed). See Figure 1, panel B.

Following failure, the stimulus was reset to the center of the screen, and the lambda parameter was reset to 50% of the value achieved at the time of the crash - allowing participants to ‘reset’ and build back up to a higher difficulty. This process was repeated 10 times. We estimated each participant’s overall motor stability threshold (MST) by calculating the average lambda values reached over the final three calibration trials.

##### 2.3.1.2 Continuous Performance Phase

In the continuous performance phase, participants performed the same task as in the calibration phase. However, in this phase the difficulty level was held to just 30% of the participant’s individually estimated MST, and the trial duration was fixed at 10 minutes. See Figure 1, panel C. To evaluate task compliance, we examined the participants’ mean position. Participants were able to maintain the position of the stimulus near the center of the screen, with an average distance of -0.156 +/- 0.168 DVA across all participants. See Figure 1, panel D.

#### 2.3.2 Flanker Task

Participants performed a modified version of a flanker paradigm (Botvinick et al., 1999; Colcombe et al., 2004) in which they were asked to respond to a central target flanked by an array of distractors. Each trial presented one of three trial types: congruent, where the flanking stimuli matched the central target (e.g., <<<<<); incongruent, where the flankers opposed the central target (e.g., <<><<); or neutral, where the flankers provided no directional information (e.g., - - < - -). Trial types were presented in equal proportions and were first-order counterbalanced to control for sequential effects.

The task consisted of a practice block of 30 trials with feedback. Each trial began with a fixation cross displayed for 500 ms, followed by the target stimulus. The inter-trial interval (ITI) averaged 1.16 seconds; mean total task duration was 618 seconds. Participants responded using a standard keyboard, pressing the "C" or "M" keys to indicate left or right central arrow directions, respectively. They were instructed to respond as quickly and accurately as possible. Participants were required to achieve at least 80% accuracy in the practice block to move on to the test phase. The test phase consisted of three blocks of 120 trials each, for a total of 360 trials. Participants were required to achieve at least 80% accuracy across all test trials to be included in analyses. 9 participants did not meet this minimum criteria.

#### 2.3.3 Woodcock Johnson Tests of Cognitive Abilities and Tests of Achievement (WJ)

Participants were administered a subset of the Woodcock Johnson Tests of Cognitive Abilities and Tests of Achievement (Schrank & Wendling, 2018) during in-person testing conducted by research staff under the supervision of the study neuropsychologist. Tests were administered according to the standardized guidelines and data were entered into the publisher’s scoring program to generate composite scores used in this analysis. Brief Intellectual Ability (BIA) is an age-normalized composite score derived from the Oral Vocabulary, Number Series, and Verbal Attention subtests. Brief Achievement (ACHBRF) is an age-normalized composite score derived from the Letter-Word Identification, Applied Problems, and Spelling subtests. Published reliability for the BIA and ACHBRF are .92 to .95 and .96 to .97, respectively, across our analysis age range (Mcgrew et al., 2014).

#### 2.3.4 Cardiorespiratory Fitness Assessment (VO2max)

VO2max was estimated using the Parvo Medics True One 2400 Metabolic Measurement System (Crouter et al., 2006) which controlled a recumbent cycle ergometer. Participants exercised at a linearly increasing workload while their heart rate, exhaled CO2, and O2 were analyzed. The assessment was terminated when users met >= 90% of their age-related heart rate maximum (220-age) and a respiratory exchange ratio (CO2:O2 ratio; RER) >= 1.02, or voluntarily terminated the session.

### 2.4 Data analyses

#### 2.4.1 cpCST Metrics

##### 2.4.1.1 Preprocessing

Raw stimulus coordinate data were preprocessed to correct for deviations caused by a crash during the continuous phase (n = 13 crashes). Crashes, identified as stimulus eccentricity exceeding ±80% of the distance from the center to the edge of the screen were removed, and the removed data were reconstructed using piecewise polynomial interpolation (PCHIP) to ensure smooth continuity. Participants with two or more crashes were classified as outliers and removed (n = 5).

##### 2.4.1.2 Instantaneous Reaction Time (iRT)

To quantify temporal responsiveness during task performance, we computed an Instantaneous Reaction Time (iRT) measure using dynamic time warping (DTW). This approach captured continuous time-varying latencies between stimulus and response movements by analyzing the x-coordinate (time) position vectors of both the stimulus object and user positions. The DTW algorithm identified the optimal alignment between these time series, producing a warp path representing temporal correspondence (See Figure 2). By multiplying x-coordinate distances by the sampling rate, we derived latency estimates for each timepoint, providing a highly granular measure of response latency. iRT computations were performed using custom Julia scripts and the DynamicAxisWarping.jl package (Carlson, 2020).

**Figure 2.**
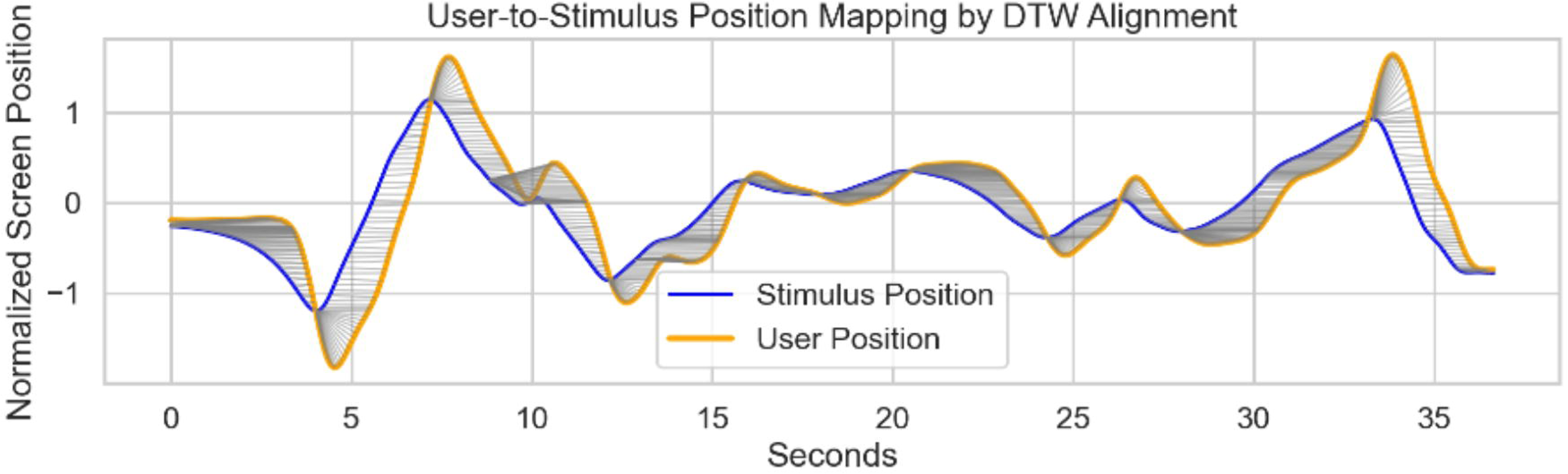
Illustration of the dynamic time warp (DTW) approach used to calculate the instantaneous reaction time (iRT) metric. The Z-scored positional timeseries for the stimulus position (blue) and the user position (orange) during the first 36 seconds of a participant’s cpCST task performance are plotted above. User position timeseries was mirrored to align with the direction of the stimulus position timeseries. DTW was then applied to find the best fit transform between the user and stimulus position. The pointwise mapping of this transform between user and stimulus position are shown as light grey lines connecting corresponding points on the blue and orange lines. iRT at each timepoint is represented by the distance in time (x axis) between the corresponding points on each line. Figure 2. Alt Text: A wide overlaid line plot titled “User-to-Stimulus Position Mapping by DTW Alignment,” showing time in seconds (0–38) on the x-axis and normalized screen position (–1.5 to 1.5) on the y-axis. A blue curve (“Stimulus Position”) and an orange curve (“User Position”) oscillate together, with numerous fine grey lines connecting matching points to illustrate the dynamic time warping alignment; a legend labels the two curves.

We then computed the mean iRT for each participant, and forwarded these to subsequent analyses.

#### 2.4.2 Flanker Metrics

Accuracy and reaction time was recorded for each trial. Participants with accuracy below 80% across all trial conditions were classified as outliers and removed (n = 15). For each participant, incorrect responses were removed from further analysis. For correct trials, anticipatory RTs, defined as RTs faster than 200ms, as well as RTs more than 2.5 SD longer than the participant’s mean were also removed from further analysis.

We computed the following metrics: mean reaction time for congruent (conRT) and incongruent (incRT) trials, and the standard flanker congruency effect (I-C; incongruent RT - congruent RT). These values were then forwarded for additional analysis.

#### 2.4.3 Reliability in cpCST and Flanker

We computed split-half reliability estimates for both the cpCST iRT and the flanker task response times (conRT, incRT, and I-C). To estimate split-half reliability and generate population-level confidence intervals, we used a bootstrap procedure (Efron, 1992). In each bootstrap iteration, participants were sampled with replacement, and split-half reliability was computed using the permutation procedure, below.

Split-half reliability In each iteration, trials were randomly permuted and split into two halves. The aggregated mean was computed for each half, and the Pearson correlation between half-scores was calculated. The Spearman–Brown prophecy formula (Brown, 1910; Spearman, 1910) was applied to correct the correlation, providing an estimate of full-test reliability. This process was repeated 1,000 times, and the average split-half reliability was reported. The split-half approach provides an index of internal consistency by estimating how well two randomly chosen halves of the test relate to each other, scaled to reflect full-test reliability.

The resulting distribution of bootstrap estimates was used to derive 95% confidence intervals (2.5th and 97.5th percentiles).

All reliability estimates were computed using custom Python code, with bootstrap iterations parallelized using Joblib for computational efficiency. Random seeds were fixed to ensure reproducibility.

#### 2.4.4 Temporal Efficiency in cpCST and Flanker

##### 2.4.4.1 Stability curves

To evaluate the temporal efficiency of each task metric, we assessed how well early portions of the task captured participants’ overall response time (RT) profiles. For each participant, we computed the mean RT separately for each task and condition using only the first n minutes of task data (e.g., first 1, 2, 3, … 9 minutes). We then correlated these truncated means with the corresponding means computed using the full duration of the corresponding task. This yielded a curve of similarity (Pearson’s r) as a function of data collection time, providing an estimate of how quickly stable RT estimates emerge for cpCST iRT and flanker-based RT metrics.

##### 2.4.4.2 Comparison of stability curves

Statistical comparison between task stability curves for cpCST and flanker trial types was performed using Steiger’s Z-test for dependent correlations with one variable in common (Steiger, 1980). For each time point (1, 2, 3, … 9 minutes), we compared the correlation between the truncated and full dataset for the cpCST iRT against the corresponding correlation for each flanker task condition. This approach appropriately accounts for the repeated measures nature of the comparison, estimating the covariance between correlations and compensating for the correlation between the truncated measures (cpCST and flanker). This provides a more conservative and accurate assessment than treating the correlations as independent (Steiger, 1980). A significant Z-statistic indicates that one task achieves temporal stability more efficiently than the other at that specific time point.

#### 2.4.5 Predictive validity

To evaluate the predictive validity of the cpCST’s instantaneous reaction time (iRT), we conducted a series of regression analyses. Specifically, we examined whether the participants’ iRT could predict performance on proximal measures (flanker task outcomes), distal measures (Woodcock-Johnson Cognition and Achievement composite scores), and a measure of central nervous system health and plasticity (VO2max). For each outcome variable, separate regression models were fitted using the mean iRT from the cpCST. We further explored the role of age, repeating these regression analyses both with and without age as a covariate in the models.

## 3 Results

Participants (N = 166) ranged in age from 18 to 76 years (M = 51.61, SD = 16.36) and reported 12 to 20 years of formal education (M = 15.81, SD = 2.11). The sample was 66% female (n = 110) and 34% male (n = 56). In terms of race, 81% identified as White (n = 134), 10% as Black or African American (n = 16), 5% as Asian (n = 8), 2% as American Indian or Alaska Native (n = 3), and 3% as multiracial (n = 5). Regarding ethnicity, 86% were Not Hispanic or Latino (n = 143), 13% were Hispanic or Latino (n = 22), and 0.6% preferred not to answer (n = 1).

### 3.1 Reliability of cpCST iRT and Flanker Outcomes

For the cpCST, the bootstrap-based estimate of population split-half reliability was high (r = .9993; 95% CI [0.999, 1.0]). Split-half reliabilities were also strong for flanker conRT(0.9846; 95% CI: [0.9824 - 0.9868]) and incRT (r = 0.9752; 95% CI: [0.9725 - 0.9780]. Although the split-half reliability for cpCST iRT was statistically greater than the flanker conRT and incRT (p. < 0.05), the absolute difference (e.g. 0.9993 vs 0.9842) is not likely meaningful.

We also assessed the reliability of the standard flanker congruency effect (I-C). The bootstrap-based reliability estimate for this difference score was significantly lower than the cpCST iRT or flanker conRT and incRTs (r = 0.8596; 95% CI: [0.8389 - 0.8805]).

### 3.2 Age and Sex Differences in cpCST and Flanker Measures

To examine potential individual differences in the primary outcome measures, we performed a series of multiple regression analyses examining the impact of age on the cpCST and flanker measures. See Figure 3 for scatterplots of flanker RTs, cpCST motor stability threshold (MST) and instantaneous reaction time (iRT) as a function of age.

**Figure 3.**
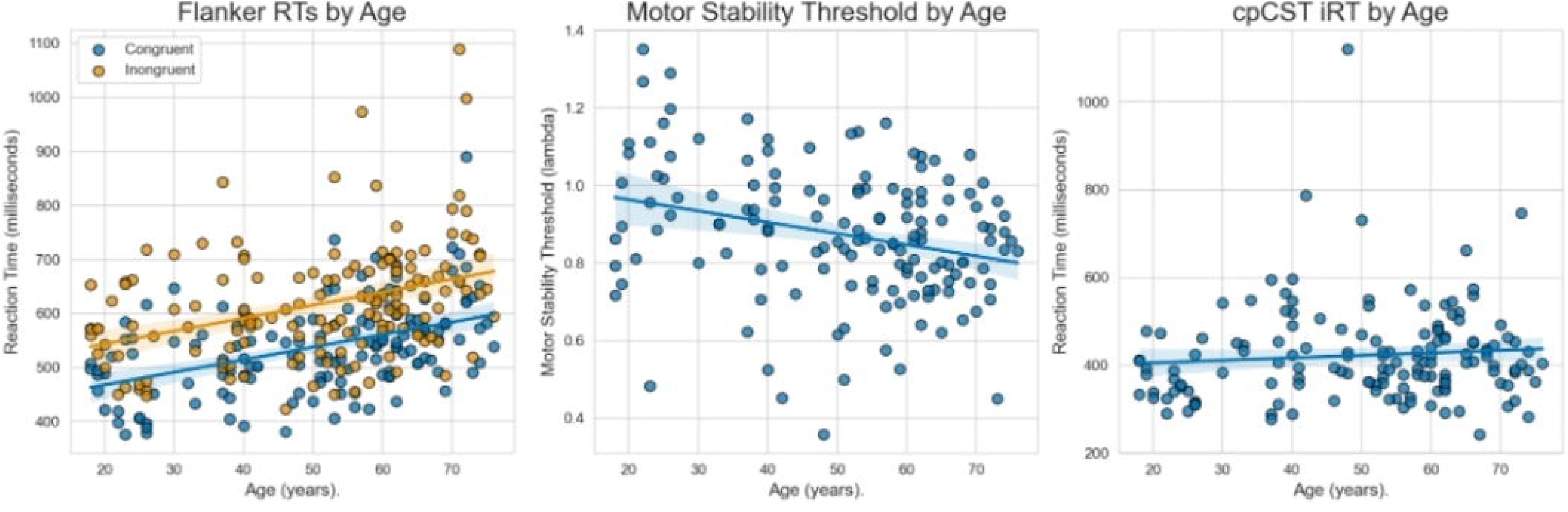
shows the association between age and the flanker task’s Congruent and Incongruent RTs, the cpCST’s Motor Stability Threshold, and cpCST iRT. Both flanker RTs and MST were significantly associated with age. The mean iRT was not significantly associated with age, suggesting that the calibration based on MST was successful. Figure 3 Alt Text: 1. **Left (Age by Flanker Grand Mean RT):** Congruent and Incongruent flanker reaction times as a function of age. Ages from 18 to 75 on x; for both Congruent and Incongruent trial types, there is a noticeable positive trend, with RTs increasing from around 450 ms in younger adults to about 600 ms or more in older adults. 2. **Middle (Age by Motor Stability Threshold):** Ages on x; calibrated stability threshold (lambda, from 0.35 to 1.35) on y. Points show a clear negative slope: younger participants tend to tolerate higher lambda values, whereas older participants cluster around lower thresholds. 3. **Right (Age by iRT):** Ages from 18 to 75 on x; mean instantaneous reaction time (iRT) in seconds (from 0.25 to 1.1 seconds) on y. Points are widely scattered around 0.4 seconds with a very slight upward trend, indicating minimal increase in mean iRT with age.

#### 3.2.1 cpCST Measures

Age significantly predicted the Motor Stability Threshold (MST; B = -0.0025, p < .001; F(3, 142) = 57.17, p < .001, R² = .55). For instantaneous reaction time (iRT), age was not a significant predictor (B = 0.0006, p = .257; F(2, 143) = 3.36, p = .038, R² = .05). These results indicate that the calibration procedure effectively adjusted for well-documented age-related slowing throughout adulthood.

#### 3.2.2 Flanker Measures

Age significantly predicted conRT (B = 2.35, p < .001; F(2, 143) = 19.99, p < .001, R² = .22) and incRT (B = 2.43, p < .001; F(2, 143) = 11.94, p < .001, R² = .14). However, age was not a significant predictor of the I-C congruency effect (B = 0.08, p = .755; F(2, 143) = 0.07, p = .933, R² = .001).

### 3.3 cpCST iRT Predictive Validity

To examine the relationship between cpCST iRT and each of our predicted metrics (Flanker, WJ, and VO2max), we conducted a series of linear regression analyses, both with and without age as a covariate.

#### 3.3.1 Flanker Features

cpCST iRT significantly predicted conRT (B = 112.48, p = .049; F(2, 143) = 20.96, p < .001, R² = .23) and incRT (B = 171.91, p = .024; F(2, 143) = 14.02, p < .001, R² = .16). However, iRT did not significantly predict the I-C congruency effect (B = 59.42, p = .112; F(2, 143) = 1.33, p = .268, R² = .02). When age was included in the models, iRT continued to significantly predict conRT and incRT, while still failing to predict the I-C congruency effect.

Combined, these findings suggest that the cpCST iRT is more closely associated with the response generation aspects of flanker task performance rather than the inhibition of conflicting responses.

#### 3.3.2 WJ Brief Intellectual Ability and WJ Brief Achievement

iRT significantly predicted WJ Brief Intellectual Ability (BIA; B = -1.79, p = .004; F(2, 143) = 8.98, p < .001, R² = .11) and WJ Brief Achievement (ACH; B = -1.42, p = .011; F(2, 134) = 8.74, p < .001, R² = .12). Faster iRT was associated with higher ability and achievement scores. Including age in the models did not eliminate these associations, suggesting that the relationships between iRT and the WJ outcome measures were not driven by age.

#### 3.3.3 VO2max

Mean iRT significantly predicted VO2max (B = -9.94, p = .010; F(2, 143) = 34.51, p < .001, R² = .33). When controlling for age, iRT remained a significant predictor of VO2max, demonstrating an association of faster reaction time speed with better aerobic capacity, beyond age-related effects.

### 3.4 Temporal Efficiency

The statistical comparison of task stability curves described how well early segments of the task captured participants’ full-task response time (RT) characterizations. The correlation for each mean cumulative (1-9) minute segment of each task’s RT features are plotted below in Figure 4. Even 1 minute of iRT data shows very good correlation with the full 10 minute assessment (r = 0.87), and by the second minute the correlation with the full sample reached r = 0.94. The flanker Congruent and Incongruent RTs also performed well, though somewhat less well than the iRT. The I-C congruency contrast performed less well than either the iRT or the base flanker features. Locations denoted by a dot on each line show where the correlations for the flanker-based RT features are significantly lower than iRT, using Steiger’s Z-test for dependent correlations with one variable in common (Steiger, 1980).

**Figure 4.**
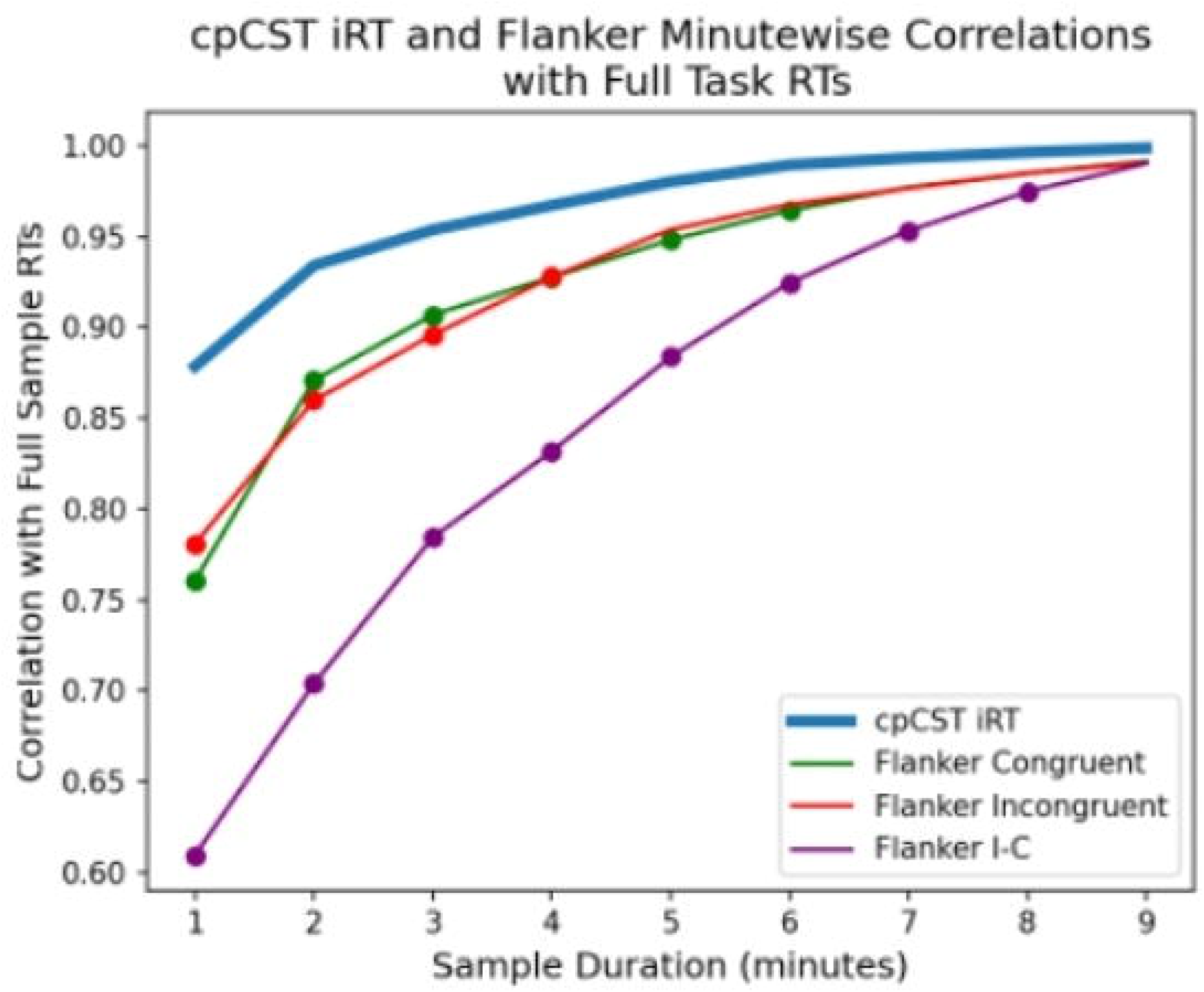
Correlation of cpCST and flanker task mean reaction times at each minute of the task compared to full task performance. Note that cpCST iRT correlation (blue line) is highest over all durations of the task, while the standard flanker congruency effect (I-C; purple line) is lowest. Figure 4 Alt Text: A line chart titled “cpCST IRT and Flanker Minute-wise Correlations with Full Task RTs.” The x-axis is Sample Duration (minutes) from 1 to 9; the y-axis is correlation with full sample RTs (0.5 to 1.0). Five series with legend labels: • **cpCST IRT (blue):** starts around 0.88 at 1 min and rises smoothly to nearly 1.00 by 9 min. • **Flanker Neutral (orange):** from 0.78 at 1 min to 0.99 at 9 min. • **Flanker Congruent (green):** from 0.76 to 0.99 over the same span. • **Flanker Incongruent (red):** from 0.78 to 0.99. • **Flanker I–C (purple):** lower starting around 0.56, climbing steadily to 0.97. All lines slope upward, with cpCST IRT highest at each minute and Flanker I–C lowest.

**Figure 5.**
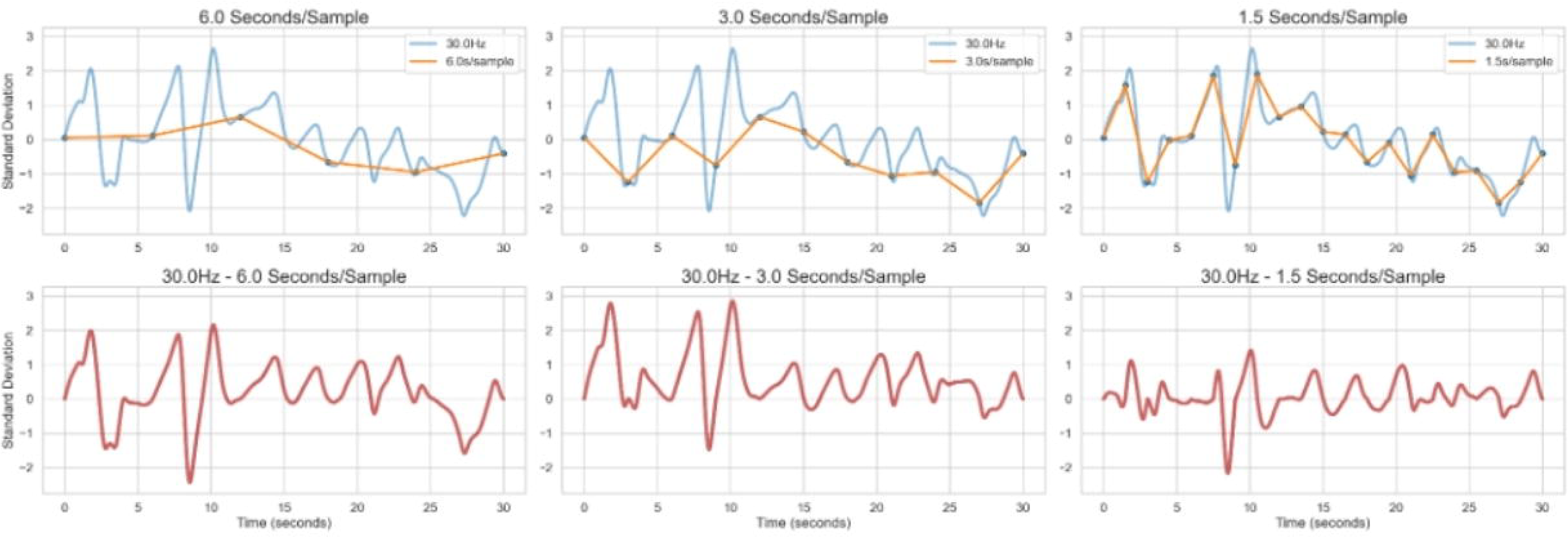
Illustration of how dense sampling of behavior may more efficiently characterize an individual’s attentional functioning. The top row shows the Z-scaled spatial path of a participant’s stimulus object (SO) over time (blue line). The SO drifts away from center (zero on the Y axis) and is subsequently returned to center; behavior is sampled at 30Hz and demonstrates a rich pattern of change over time. The top row also shows that trajectory of behavior, but sampled at rates in the range of standard discrete RT paradigms such as the PVT and Conners CPT (orange lines; 6.0, 3, and 1.5 seconds, left to right), demonstrating a much simplified pattern of apparent behavior. In the bottom row, we subtract the simulated discrete RT behavior from the same behavior as sampled at 30Hz in the cpCST. Examining these plots, it is evident that much of the behavioral variation evident in the cpCST remains undescribed if sampled in the typical discrete RT task sampling temporal regimes. This is most evident when sampled every 6 seconds. However, even when the sampling rate is increased to 1.5 seconds and the orange line more closely matches the 30 Hz blue line, the subtraction plotted in red reveals nontrivial variation. In all plots periods of a participant’s behavioral variation greater than 2 standard deviations would remain unaccounted under traditional discrete RT paradigms. This improved precision in assessment may help to explain the cpCST’s high split-half reliability and temporal efficiency. Figure 5 Alt Text: Six-panel figure showing how downsampling affects the standard-deviation trace of a 30 Hz signal over 30 seconds. All panels share “Time (seconds)” on the x-axis (0 to 30) and “Standard Deviation” on the y-axis (approximately 2.5 to +3). **Top row (overlays):** 1. **6.0 Seconds/Sample:** A blue continuous line (“30.0 Hz”) with pronounced peaks around 2 seconds, 8 seconds, and 11 seconds, overlaid by a smoother orange line with markers (“6.0 seconds per sample”) that roughly captures the overall shape of the blue line at coarse intervals. 2. **3.0 Seconds/Sample:** An identical blue trace, overlaid by an orange line (“3.0 seconds per sample”) that more closely follows the blue pattern but with fewer data points. 3. **1.5 Seconds/Sample:** An identical blue line, now with an orange line (“1.5 second/sample”) that nearly coincides with the blue line, but fails to capture finer features of the blue line. Each top panel includes a legend in the upper right identifying the blue and orange series. **Bottom row (differences):** 4. **30.0 Hzt to 6.0 Seconds/Sample:** A single red line showing the point-by-point difference between the 30 Hz and 6 seconds/sample traces, with residual peaks up to ±2.5 SD. 5. **30.0 Hz to 3.0 Seconds/Sample:** A red residual trace with peaks up to about ±2.7 SD, reflecting smaller downsampling error. 6. **30.0 Hz to 1.5 Seconds/Sample:** A red residual trace showing a smaller, but still meaningful (spanning ±1.2 SD) difference at finer sampling. Panels are arranged in a 2×3 grid with their titles centered above each subplot.

## 4 Discussion

We introduced the cpCST, a novel test of sustained attention offering high temporal precision across the lifespan. The cpCST provides reliable, valid, and temporally efficient measurement of attentional control - offering a valuable complement to traditional attention assessment paradigms. Here we summarize key methodological innovations and psychometric properties, followed by implications for future research and clinical applications.

### 4.1 Methodological Innovations

The cpCST incorporates three central methodological innovations.

High-density behavioral sampling (30 Hz) captures behavior at a granularity not possible with traditional discrete-response continuous performance tasks, which as noted above typically sample at rates of 0.1 to 1 Hz (every 1-10 seconds; cf (Basner & Dinges, 2011; C. Conners et al., 2000; C. K. Conners et al., 2003; Dinges & Powell, 1985; Homack & Riccio, 2006)). The enhanced temporal resolution provides data ideally suited to integrate with other data modalities such as EEG and physiological metrics - allowing sophisticated analyses of attentional stability and variability.

We also created a novel instantaneous reaction time measure, which estimates the temporal lag between the movement of a central stimulus object and the participant’s response to adjust to that movement. This approach estimates response time with high precision, reliability, and excellent temporal efficiency.

Additionally, the cpCST utilizes a hybrid design that combines adaptive calibration and subsequent fixed-difficulty assessments. Integrating the strengths of adaptive and fixed-difficulty paradigms provides individualized task difficulty while avoiding issues common in fully adaptive methods, such as oscillatory artefacts or instability (García-Pérez, 2011; Kontsevich & Tyler, 1999). It may also obviate the need for alternative task forms across groups with disparate baseline functioning, or in highly heterogeneous samples such as in aging, developmental, or lifespan studies.

### 4.2 Psychometric Properties

The cpCST yielded high reliability estimates, with bootstrap-based split-half reliability greater than 0.999. High-density sampling and individualized calibration likely contributed to this reduced measurement error, facilitating the rapid detection of subtle individual differences (r > 0.9 after 1 minute of data). This reliability may be especially advantageous in longitudinal studies or in interventions examining modest performance changes.

Age invariance is a notable strength of the cpCST. Although motor stability thresholds (MST) and traditional reaction time measures from the flanker task exhibited expected age-related slowing, cpCST’s iRT was stable across age. By calibrating task difficulty to each individual’s sensorimotor capacity, the cpCST appeared to effectively isolate attentional control from baseline sensorimotor function. This makes the task especially suitable in lifespan cognitive assessments, circumventing the need for distinct age-specific task versions.

The cpCST also exhibited robust validity across multiple domains. Significant associations with flanker conRT and incRT suggest convergent validity. However, the lack of association with the flanker congruency effect indicates the cpCST primarily captures tonic aspects of attention (e.g., vigilance, sustained focus) rather than the application of inhibitory control processes. Furthermore, significant predictive relationships with WJ Brief Intellectual Ability, Brief Achievement, and physiological measures like VO2max underscore the cpCST’s potential in describing fundamental cognitive and physiological features relevant to attentional control (Colcombe & Kramer, 2003; Colcombe et al., 2004; Kramer & Colcombe et al., 2018; Posner & Petersen, 1990).

The temporal efficiency of the cpCST was also notable. Within 2 minutes, the cpCST iRT exceeded an r = 0.9 correlation with full task performance. By comparison, the flanker trial types needed roughly 5 minutes of data to reach this level of association with the full flanker sample.This suggests the potential for cpCST to reduce task administration time without significant loss of information.

### 4.3 Implications and Future Directions

The central features of the cpCST position it as a promising tool for research and clinical settings. Its high temporal resolution enables tighter integration with physiological measures (e.g. EEG, fMRI, heart rate, skin conductance), facilitating exploration of neural mechanisms underlying moment-to-moment attention variability, and “brain-body” interactions. Additionally, its individualized calibration method is likely to prove valuable in heterogeneous clinical populations or lifespan studies, as it reduces confounds related to sensorimotor speed differences or ceiling/floor effects.

The task’s temporal efficiency and straightforward administration suggest suitability for large-scale assessments, longitudinal monitoring, and remote or mobile implementations. Future studies should explicitly evaluate cpCST’s sensitivity to attentional changes resulting from interventions (e.g., sleep deprivation, stimulant medication, cognitive training) and establish its utility in diverse clinical populations (e.g., ADHD, TBI, MCI). Additionally, as illustrated in Figure 6, the high density sampling may allow detection of subtle behavioral dynamics not captured with discrete response paradigms - which may not only contribute to the cpCST’s relatively high temporal efficiency and reliability, but also allow for new insights into attentional dynamics.

Finally, the rich, high-density behavioral data generated by the cpCST is well-suited for computational modeling approaches, such as drift diffusion models or Bayesian frameworks. Future work could leverage these modeling techniques to better characterize the attentional process dynamics captured by the cpCST.

### 4.4 Limitations

Several limitations should be acknowledged. Although validity and split-half reliability were established, the cpCST’s sensitivity to intervention-induced attentional changes remains to be validated. Our current age range (18-79 years) is substantial, yet the efficacy and validity in younger and older extremes require further examination. Likewise, this is a community-based normative sample and psychometric properties should be evaluated across different clinical populations. Given the cross-sectional nature of our sample, we can only establish internal reliability through bootstrap methods. Future work is needed to examine test-retest reliability under frameworks like the intraclass correlation coefficient (ICC; (Shrout & Fleiss, 1979; Zuo & Xing, 2014)).

Finally, while high-density behavioral sampling offers analytical richness, the relative complexity of calculating iRT using dynamic time warping (DTW) may present obstacles to widespread adoption. To address this, we will provide streamlined and containerized analysis pipelines on GitHub. Developing accessible pipelines and normative databases will be essential for broader clinical adoption and research utilization.

### 4.5 Conclusion

The Continuous Performance Critical Stability Task introduces methodological advances in the assessment of sustained attention. Its exceptional reliability, age invariance, predictive validity, and temporal efficiency address limitations in existing attentional measures. Future validation efforts integrating physiological measures, computational modeling, and diverse clinical applications will further establish cpCST’s utility as an essential tool for attention research and assessment.

## 5 Data Availability Statement

The datasets generated for this study can be found in the NKI-Rockland Sample data resource https://rocklandsample.org/. Access to raw data requires completion of a data use agreement. Analysis datasets used for the current study are available from the corresponding author upon reasonable request.

## 6 Author contributions

A.M.B. framed portions of the research question, oversaw and contributed significant portions of the analysis, and drafted significant portions of the article; D.G.B, K.G., O.R. and E.G. contributed to data management and analysis and drafted portions of the article; M.M. provided general guidance and edited portions of the article; S.C. oversaw the project, framed the research question, oversaw and contributed significant portions of the analysis, and drafted significant portions of the article.

## 7 Funding

This work was supported by the National Institute of Mental Health (R01MH124045; *The NKI Rockland Sample II: An Open Resource of Multimodal Brain, Physiology & Behavior Data from a Community Lifespan Sample*). Additional support was provided by the New York State Office of Mental Health and the Nathan Kline Institute institutional core services. Author salaries were supported by New York State (A.M.B., S.C., M.M., E.G.) and by the NIH grant R01MH124045 (D.G.B., O.R., K.G.).

## 8 Acknowledgements

We acknowledge the important support of the entire Nathan Kline Institute – Rockland Sample team of investigators, research support staff, NKI Scholars and, most importantly, the community member participants who volunteer time and energy to advance scientific knowledge and benefit others.

## 9 Conflict of Interest

The authors declare that the research was conducted in the absence of any commercial or financial relationships that could be construed as a potential conflict of interest.

## 10 Generative AI Statement

The authors declare that no Generative AI was used in the creation of this manuscript.

## Notes

https://rocklandsample.org/

### Competing Interest Statement

The authors have declared no competing interest.

